# Functionally redundant forms of extended-spectrum beta-lactamases and aminoglycoside-modifying enzymes drive the evolution of two distinct multidrug resistance gene clusters in clinical populations of EXPEC

**DOI:** 10.1101/367938

**Authors:** Jay W. Kim, Portia Mira, Patricia P. Chan, Todd M. Lowe, Miriam Barlow, Manel Camps

## Abstract

We evaluate the distribution of genetic markers for antibiotic resistance in 276 genomic sequences of Extraintestinal Pathogenic *E. coli* from two hospitals on the U.S. West coast. Plasmid-borne genes encoding drug-inactivating enzymes dominate the distribution of aminoglycoside and *β*-lactam resistance markers. These genes can be assigned based on their distribution to two mutually exclusive complementarity groups (CGs: CG1 and CG2) with each displaying genetic linkage and minimal functional overlap. CG1 includes genes encoding OXA-1 and AAC(6’)-Ib-cr, frequently also CTX-M-15, and sometimes AAC(3)-IIe.2, a variant of AAC(3)-IIe; CG2 includes AAC(3)-IId tightly linked to TEM-1, and occasionally also to genes encoding CTX-M-14-like *β*-lactamases. This binary distribution of aminoglycoside and *β*-lactamase resistance genes suggests a convergence between two different evolutionary solutions, and results in a ubiquitous functional redundancy in the clinical populations. CG1 and CG2 are largely carried in IncF plasmids, of which we distinguish seven classes based on Rpt-A1 sequence homology. Both CG1 and CG2 genes are found in two different IncF plasmid classes, demonstrating their pervasive mobility across plasmid backbones. Different CG genes and IncF plasmid classes are found in a wide range of MLSTs, highlighting the prevalence of horizontal gene transfer. We also identify at least five clonally expanding MLSTs, which represent high-risk clones: ST131, ST95, ST73, ST127, and ST69. The identification of clonally-expanding types, the discovery of CGs that are ubiquitously spread in diverse clinical strains, and the functional redundancy that these two groups represent have significant implications for monitoring and controlling the spread of resistance.

## Introduction

Antibiotic resistance represents a growing threat to public health, increasing medical costs worldwide. Particularly, the spread of multidrug-resistant *E. coli* clones is a serious public health concern because of its symbiotic relationship with humans and animals (Li et al. 2007; Peirano et al. 2012), and because of its frequent implication in opportunistic infection of its hosts, causing urinary tract infections, sepsis and wound infections both in clinical and community-associated settings (Robicsek et al. 2006; Price et al. 2013; Yano et al. 2013). The spread of resistance is further accelerated by incorrect or unnecessary antibiotic treatments (Ventola 2015; Burnham et al. 2017). Thus, understanding the mechanisms driving the spread of resistance and improving the accuracy of therapeutic decisions through point-of-care diagnostics should be of great help to both slow down the development of antibiotic resistance and to reduce the costs associated with antibiotic-resistant infections.

Here, we evaluate the distribution of known antibiotic resistance genes in 276 genomic sequences of Extraintestinal Pathogenic *E. coli* (ExPEC) isolated in two West Coast, U.S. hospitals: University of Washington (UW) in Seattle, Washington (Salipante et al. 2015), and the Dignity Health Mercy Medical Center (DHMMC), in Merced, California. We focus on resistance to the three classes of antibiotics most frequently used for the treatment of ExPEC infections and of enterobacterial infections in general, namely aminoglycosides, *β*-lactams, and fluoroquinolones (Rodriguez-Bano et al. 2018) (for a list of antibiotics included in this study and their distribution in the population see **Methods)**.

Aminoglycosides are a large class of structurally related compounds that inhibit different steps of bacterial protein synthesis (Magnet and Blanchard 2005; Jana and Deb 2006; Shakil et al. 2008). Resistance can result from direct modification and inactivation of the drug molecule by aminoglycoside-modifying enzymes (Ramirez and Tolmasky 2010), methylation of the amino acyl site of 16S rRNA through 16S rRNA methyltransferases (Doi et al. 2016), and by mutations in genes encoding ribosomal proteins (e.g. *rpsL*). Additional contributors to aminoglycoside resistance include altered expression and/or mutations in outer membrane porins, two-component systems and efflux systems (Cag et al. 2016).

*β*-lactam antibiotics include penicillins, cephalosporins, monobactams and carbapenems. These drugs diffuse into the cells’ periplasm, where they bind to and inactivate penicillin-binding proteins, which are essential for cell wall maintenance, resulting in cell death (Zapun et al. 2008; Sauvage and Terrak 2016). Resistance to *β*–lactams is largely due to hydrolysis by *β*–lactamases, which in ExPEC include TEM, SHV, CTX-M, OXA, AmpC, and a variety of carbapenemases (Gutkind et al. 2013). All of these proteins have activity against penicillins. Some of them can also protect against oxyimino (3^rd^ and 4^th^ generation) cephalosporins and against monobactams, a phenotype known as Extended-Spectrum *β*–Lactamase (ESBL) resistance (Ghafourian et al. 2015). ESBL activity can be constitutive (CTX-Ms) or evolved, as is the case of mutant TEM, mutant SHVs, and some OXAs. Carbapenems have been introduced more recently for the treatment of ESBL resistance, and only carbapenemases (KPC, NDM, VIM) (Martinez-Martinez and Gonzalez-Lopez 2014; Jeon et al. 2015) and most *β*–lactamases of the OXA family exhibit activity against them (Mairi et al. 2017).

Fluoroquinolones inhibit type-II topoisomerase activity, blocking replication. Resistance frequently derives from mutations in the two main topoisomerase II-type enzymes present in *E. coli*, namely DNA gyrase and DNA topoisomerase IV. Each consists of two subunits: GyrA and GyrB for DNA gyrase and ParC and ParE for DNA topoisomerase IV. In *E. coli* and other Gram negative pathogens, DNA gyrase is the preferential target for fluoroquinolones but highly-resistant organisms tend to contain resistance-conferring mutations in both enzymes (Redgrave et al. 2014). These mutations tend to occur specifically in the quinolone resistance determining regions (QRDR) of GyrA and ParC, preventing the binding of the drug to these targets (Redgrave et al. 2014).

Thus, a considerable number of drug resistance factors have already been identified. The emerging picture is that antibiotic resistance involves sequential lines of defense targeting drug influx, accumulation, target binding, or downstream toxicity (Yelin and Kishony 2018). We are also beginning to understand the processes leading to the acquisition and spread of resistance genes in the population, but significant gaps remain due both to the complex distribution of pathogens within hospitals and across the community (Peirano et al. 2012; Kang et al. 2013), and also due to the difficulty in adequately detecting resistance genes by PCR or molecular hybridization methods because these genes can exhibit a high degree of genetic variability (Leinberger et al. 2010; Rubtsova et al. 2010). With the advent of next-generation sequencing, the application of WGS technologies in clinical diagnostics and research has been a fast-growing trend, particularly in tracking hospital outbreaks of bacterial pathogens (Snitkin et al. 2012; Luo et al. 2014; Fitzpatrick et al. 2016), and predicting drug resistance based on bacterial sequence rather than with phenotypic assays (Gordon et al. 2014; Coll et al. 2015; Walker et al. 2015).

Plasmids are autonomously replicating units that greatly facilitate the spread of antibiotic resistance (Carattoli 2013; Brolund and Sandegren 2016). Most aminoglycoside-modifying enzymes and *β*–lactamases are carried in plasmids (Banerjee and Johnson 2013; Brolund and Sandegren 2016). While the main mutations conferring fluoroquinolone resistance are chromosomal, plasmid-borne proteins of the pentapeptide repeat family encoded by quinolone resistance (*qnr*) genes can also confer moderate fluoroquinolone resistance by protecting DNA gyrase and topoisomerase IV (Rodriguez-Martinez et al. 2016; Yanat et al. 2017). Finally, the aminoglycoside acetyltransferase AAC(6’)-Ib-cr can acetylate ciprofloxacin and norfloxacin, thus contributing to resistance (Briales et al. 2012).

Plasmids carry mobile elements including insertion sequences (IS), integrons, and transposition units (reviewed in (Partridge 2011; Guedon et al. 2017)) that facilitate the capture of resistance genes from a shared environmental pool (Crofts et al. 2017). Each particular resistance gene is generally immediately associated with a specific mobile element, possibly reflecting individual gene capture processes (Partridge 2011). Through recombination, these mobile elements in turn also facilitate the acquisition of additional antibiotic resistance genes through a stochastic process known as *genetic capitalism*, which favors the acquisition of additional antibiotic resistance genes by strains already exhibiting antibiotic resistance (Canton et al. 2003; Canton and Ruiz-Garbajosa 2011) and also the clustering together of these antibiotic resistance genes in large multi-resistance regions (MRRs) (Baquero 2004; Partridge et al. 2011). Plasmids also facilitate the spread of resistance through conjugation, a contact-dependent and energy-driven process mediated by a molecular machinery that is related to type IV secretion systems (Ilangovan et al. 2015; Banuelos-Vazquez et al. 2017). This means that plasmids can transfer across unrelated bacterial strains, spreading antibiotic resistance genes in the process (Shin et al. 2012; Hu et al. 2014).

The minimal portion of a plasmid replicating with the characteristic copy number of the parent is known as a replicon. Replicons are classified into incompatibility (Inc) groups based on their ability to be propagated stably in the same cell (Novick 1987). A total of twenty-seven Inc groups are known in *Enterobacteriaceae*, although their representation varies substantially in naturally occurring bacteria. IncF plasmids are one of the most widely diffused plasmid families in clinically relevant *Enterobacteriaceae*,(Johnson et al. 2007) and have played a major role in the dissemination of bla_CTX-M-15_, and bla_CTX-M-14_ through IS*Ecp1* capture and mobilization and through their association with the high-risk strain ST131 (Nicolas-Chanoine et al. 2014). IncF plasmids are unusual in that they frequently contain multiple replicons, possibly resulting from the recombination of different plasmids (Villa et al. 2010). In addition, IncF plasmids have an abundance of mobile elements, which also facilitate genetic exchange. As a result, IncF plasmids show a very high level of recombination, with widespread deletions and truncations, and highly variable modules associated with virulence and biochemical pathways (Partridge et al. 2011; Li et al. 2015).

The adaptive success of *high-risk clones* (host strains) often contributes to the spread of plasmid-mediated antibiotic resistance. Sequence type 131 (ST131) is a case in point, evolving resistance to fluoroquinolones through *gyrA* and *parC* mutations (Cagnacci et al. 2008; Johnson et al. 2013), acquiring bla_TEM-1_ and bla_OXA-1_ IncFII plasmids, with the subsequent capture of ESBL β–lactamases (Woodford et al. 2011; Olesen et al. 2013), and often also capturing aminoglycoside resistance genes (Li et al. 2015).

Here, we conducted a large-scale genomic study using data derived from ExPEC isolates from two U. S. West Coast hospitals to identify the distribution of known genetic antibiotic resistance genes in ExPEC. We found extensive genetic exchange associated with IncF plasmid recombination and widespread Horizontal Gene Transfer (HGT), allowing the co-existence of alternate determinants of *β*–lactam or aminoglycoside resistance in the population. Further, we find that alternate determinants of resistance are genetically linked, forming two complementarity groups. The identification of distinct complementarity groups has substantial implications for our understanding of how drug resistance evolves as well as for diagnosis and monitoring of antibiotic resistance.

## Results

We performed whole genome sequencing of 45 ESBL-resistant ExPEC isolates from Dignity Health Mercy Medical Center (DHMMC) and obtained 384 assembled draft genomes from a published University of Washington (UW) study (Salipante et al. 2015). Both of these sets had been tested for susceptibility against a panel of antibiotics, with some differences in panel composition between the two. From these, we selected 276 genomes (43 from DHMMC and 233 from UW) by removing longitudinal duplicates and controlling for sequencing quality (Details about the selection of clinical samples and the processing of the sequences are provided in *Methods* and *Supplemental Material*). The antibiotics considered in our study are shown in **Supplemental_Table_S1**.

### Phylogenetic structure

As an initial indication of strain diversity, we performed multilocus sequence typing (MLST) according to the Achtman classification (Maiden et al. 1998) (**Supplemental_Data_S1**). The most abundant MLSTs included ST131 (65 samples), ST95 (27 samples), ST73 (22 samples), ST127 (15 samples) and ST69 (12 samples). At the other end of the spectrum, 68 of the 276 genomes corresponded to MLSTs with 3 or less representatives. To obtain higher resolution, we generated a phylogeny based on SNP variation across the genome (**Fig. 1**).

**Figure 1.**
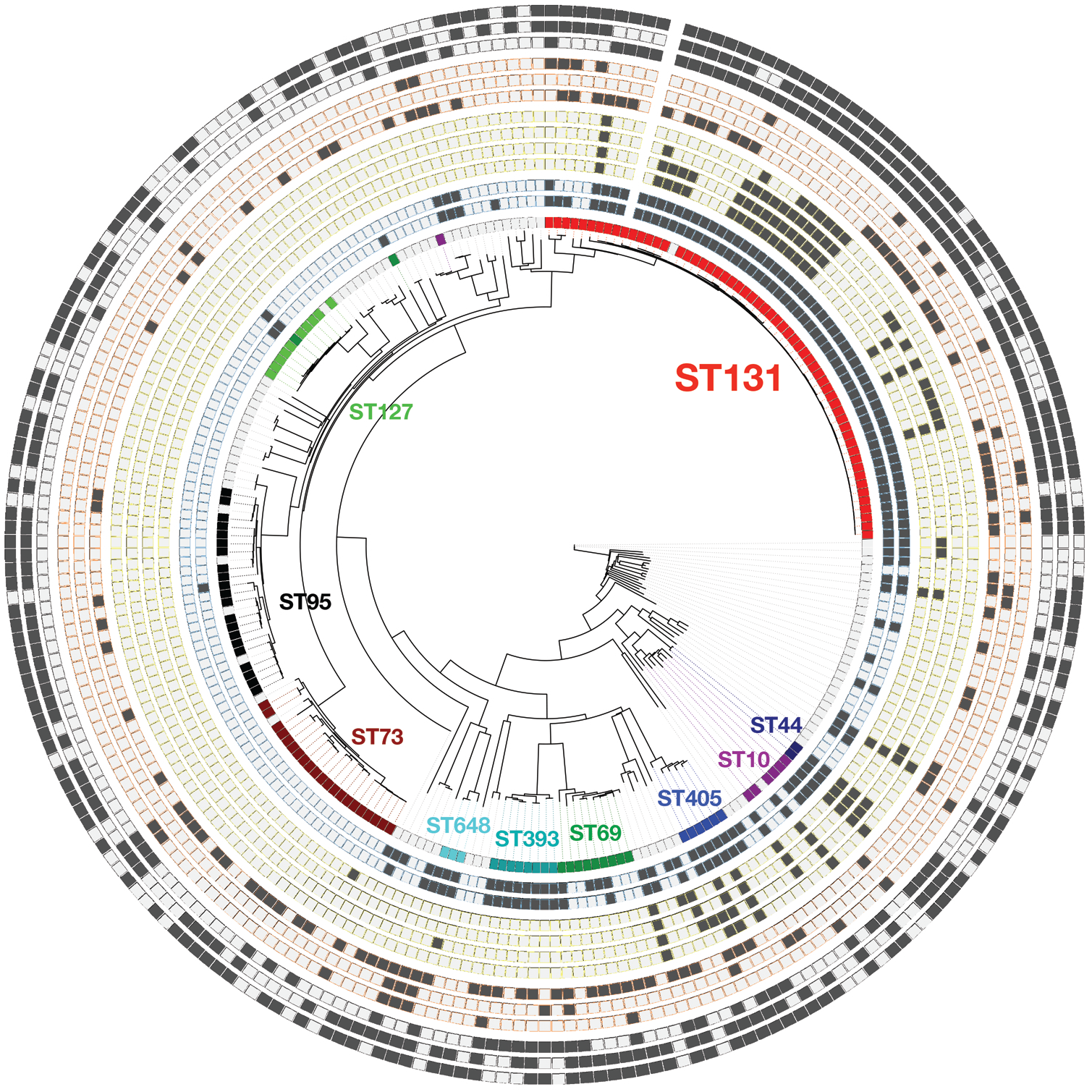
Phylogeny of 276 ExPEC strains obtained from two U.S. West Coast hospitals. Phylogenetic relationships among these strains are represented as a neighbor-joining tree computed based on pairwise comparisons of genomic SNPs. The Circos diagram surrounding the phylogeny shows, for each phylogenetic taxa, an MLST classification, presence of various resistance markers and plasmid replicons. Circular panels represent the following from the innermost tooutermost positions. **Panel 1**, MLST classification: ST131 (red), ST44, (navy), ST10 (purple), ST405 (blue), ST69 (green), ST393 (teal), ST648 (cyan), ST73 (maroon), ST95 (black), ST127 (lime). **Panels 2** and **3** (outlined in blue), fluoroquinolone resistance mutations:

GyrA-S83X and ParC-S80I, respectively. **Panels 4–8** (outlined in yellow), complementarity group 1: genes for OXA-1, AAC(6’)-Ib-cr, CTX-M-15, and AAC(3)-IIe.2, respectively. **Panels 9–11** (outlined in orange), complementarity group 2: genes for TEM-1, CTX-M-9-family and AAC(3)-IId, respectively. **Panels 12–14** (outlined in grey), IncF replicons: IncFIA, IncFIB, IncFII, respectively.

In this phylogeny, clonal expansion appears as poorly resolved clades, corresponding to the following MLSTs: ST131, ST95, ST73, ST127, and ST69. Consistent with recent expansion, these types coincide with the most well represented MLSTs. In general, the phylogeny is in good agreement with the MLST classification, with some exceptions (*Supplemental Material*).

### Selection of antibiotic resistance markers

We compiled a list of genetic resistance genes reported for ExPEC and also for other Gram negatives. This list included both genes and point mutations known to be associated with the acquisition of resistance against drugs belonging to the three antibiotic classes that are most frequently used in the clinic for the treatment of enterobacterial infection, namely *β*–lactams, aminoglycosides and fluoroquinolones (Lanza et al. 2014; Zowawi et al. 2015; Cerceo et al. 2016; Rodriguez-Bano et al. 2018).

UniProt accessions and protein sequences for all known antibiotic resistance genes considered in our study are provided in **Supplemental_Data_S2**. Details about the drugs tested and the number of clinical isolates tested for each drug are provided in *Supplemental Material* and summarized in **Supplemental_Fig_S1**. Next, we performed sequence similarity searches for these genes on the 276 ExPEC draft genomes. Resistance genes and sequence variants found among the 276 genomes are listed in **Supplemental_Data_S2**. To investigate the mobility of these genes, we also looked for the presence of known plasmid replicons, listed in **Supplemental_Data_S1** by incompatibility group (Carattoli 2011).

### Representation of resistance genes in our dataset

We found a variety of aminoglycoside modification enzymes encoded among the 276 genomes, with a clear predominance of three specific aminoglycoside acetyltransferases: AAC(6’)-Ib-cr (34 genomes, 12.7% of the total), AAC(3)-IId (28 genomes, 10.1%) and a novel variant, AAC(3)-IIe.2 (15 genomes, 5.4%), which differs from the AAC(3)-IIe reported by Ho et al. (Ho et al. 2010) only by three amino acid substitutions: F14L, Q275H and K276E. The distribution of these three aminoglycoside acetyltransferases in our dataset is shown in **Fig. 1**, panels 5, 7 and 10.

There were also 19 genes encoding the ANT(2’’)-I adenylyltransferase or variants thereof (**Supplemental_Data_S2**), and only one encoding the 16S rRNA methyltransferase, *RmtE*. The genome bearing this methyltransferase displayed resistance to both gentamicin and tobramycin and did not contain any other aminoglycoside resistance genes within our consideration, suggesting a causal link to gentamicin and to tobramycin resistance. Another genome contained a nonsynonymous *rpsL* mutation (T39I), but it was unclear whether the *rpsL* T39I mutation contributed to gentamicin and tobramycin resistance because AAC(3)-IId was also encoded in this genome. In addition, we found genes corresponding to two aminoglycoside-modifying enzymes that confer resistance to aminoglycosides not included in this study.

There were 117 TEM, 37 OXA, and 48 CTX-M *β*-lactamases encoded among the 276 genomes. Of the six different types of plasmid-encoded AmpC *β*
-lactamases that we checked, only one was found: the CMY-2 *β*-lactamase (5 genomes, **Supplemental_Data_S2**). Note that SHV-1 *β*-lactamases were not represented in any of the genomes. We did not search for carbapenemases because carbapenem resistance was almost completely absent from our clinical sample (<1% prevalence).

We also searched for polymorphisms in the *β*–lactamases. Among the 117 TEM sequences, there were only three containing ESBL mutations, two with the G238S substitution and one with the R164S mutation. Among the 37 OXA sequences, we discovered a single OXA-1 variant bearing an M102I substitution, which is in the active site and has not been previously reported. CTX-M *β*–lactamase sequences, on the other hand, exhibited much broader diversity, including CTX-M-15 (37 genomes), CTX-M-14 (6 genomes), CTX-M-27 (3 genomes) and CTX-M-65 and CTX-M-104 (1 genome each). The distribution of CTX-M-15 and of CTX-M-14-like enzymes is shown in **Fig. 1**, panels 6 and 9, respectively.

The profile of *β*–lactamases represented in our population is consistent with evidence of CTX-M displacement of other ESBL enzymes worldwide (Rodriguez-Villalobos et al. 2011), with CTXM-14 and CTX-M-15 being the predominant ones (Peirano and Pitout 2010; Hiroi et al. 2012; Doi et al. 2013). Of note, CTX-M-27 and CTX-M-15 were found together in one of our genomes. The coexistence of different CTX-M enzymes in the same sample has been reported, and has already given rise to new CTX-M variants by recombination (Sun et al. 2010; Tian et al. 2014).

Fluoroquinolone resistance generally maps to point mutations in two type-II topoisomerases, although resistance can also be plasmid-mediated (Hopkins et al. 2005; Redgrave et al. 2014). Of the 276 genomes, 47.6% had a *gyrA* mutation (translating to S83L), which is a key resistance-conferring mutation in *E. coli* and several other Gram negatives (Redgrave et al. 2014; Johnning et al. 2015).S83L was frequently accompanied by a mutation in *parC*, S80I/R, which in combination with GyrA_S83L produces high levels of fluoroquinolone resistance (Marcusson et al. 2009). The frequent co-occurrence of these two, synergistic mutations is well established (Johnning et al. 2015) and can be clearly seen in **Fig. 1**, panels 2 and 3. By contrast to *gyrA* and *parC* mutants, which were ubiquitous among fluoroquinolone-resistant samples, we only found one plasmid-mediated quinolone resistance gene.

### Correlation of genetic markers with drug resistance

In order to infer the possible selections driving the fixation of resistance genes in our clinical populations, we performed a linear regression analysis (**Supplemental_Fig_S2**), which can identify genes that contribute to prediction of resistance for a given drug, and provide a coefficient reflecting the relative contribution of each individual gene to the predictive model. Note that the coefficients for individual genes cannot be interpreted in isolation, as some of these genes are genetically linked, *i.e*. frequently co-occur in samples (**Fig. 1**).

In the case of aminoglycosides (**Supplemental_Fig_S2, 2b**), the presence of the two AAC(3)-type acetyltransferase genes correlated preferentially with gentamicin resistance, whereas AAC(6’)-Ib-cr genes showed a preferential, although weaker, correlation with tobramycin resistance. *β*–lactams and their resistance markers displayed more complex relationships (**Supplemental_Fig_S2c-h**). CMY-2, CTX-M-14 and CTX-M-15 genes all displayed strong correlations with resistance to the cephalosporins ceftazidime (3^rd^ gen), ceftriaxone (3^rd^ gen) and cefepime (4^th^ gen). Resistance to monobactams (aztreonam, **Supplemental_Fig_S2e**), however, correlated only to the CTX-M-15 and CMY-2 genes, suggesting that CTX-M-14 has little activity against monobactams. OXA-1’s apparent correlation with resistance to these antibiotics likely arose from linkage disequilibrium between the OXA-1 and CTX-M-15 genes on IncF plasmids (see discussion of complementarity groups below).

No single *β*–lactamase gene showed a predominant contribution towards prediction of resistance to ampicillin, consistent with the ability of all of these *β*–lactamases to target ampicillin and with the diversity of *β*–lactamases encoded in the 276 genomes. For unknown reasons, cefazolin showed a linear regression profile more similar to that of 3^rd^ and 4^th^-generation cephalosporins despite being a first-generation cephalosporin, and therefore a substrate for TEM-1, OXA-1, and CMY2 (Jacoby 2009; Gutkind et al. 2013).

### Linkage structure of MDR genes

To get an initial idea of antibiotic resistance gene co-occurrence in the 276 genomes, we generated a heatmap representing the strength of association corresponding to each pair (**Fig. 2)**. Aminoglycoside acetyltransferase and ESBL *β*–lactamase genes clustered to two different groups based on their strengths of association. The first group contained AAC(6’)-Ib-cr and AAC(3)-IIe.2, while the second group had AAC(3)-IId. Similarly, *β*–lactamases also clustered to two groups. CTX-M-15 and OXA-1 clustered to the first group, while TEM-1 and CTX-M-14 family or CMY-2 clustered to the second. Mutations conferring fluoroquinolone resistance (GyrA_S83L and ParC_S80I/R), by contrast, were strongly associated with both groups, although showing a slight preference for the first group (**Fig. 2**).

**Figure 2.**
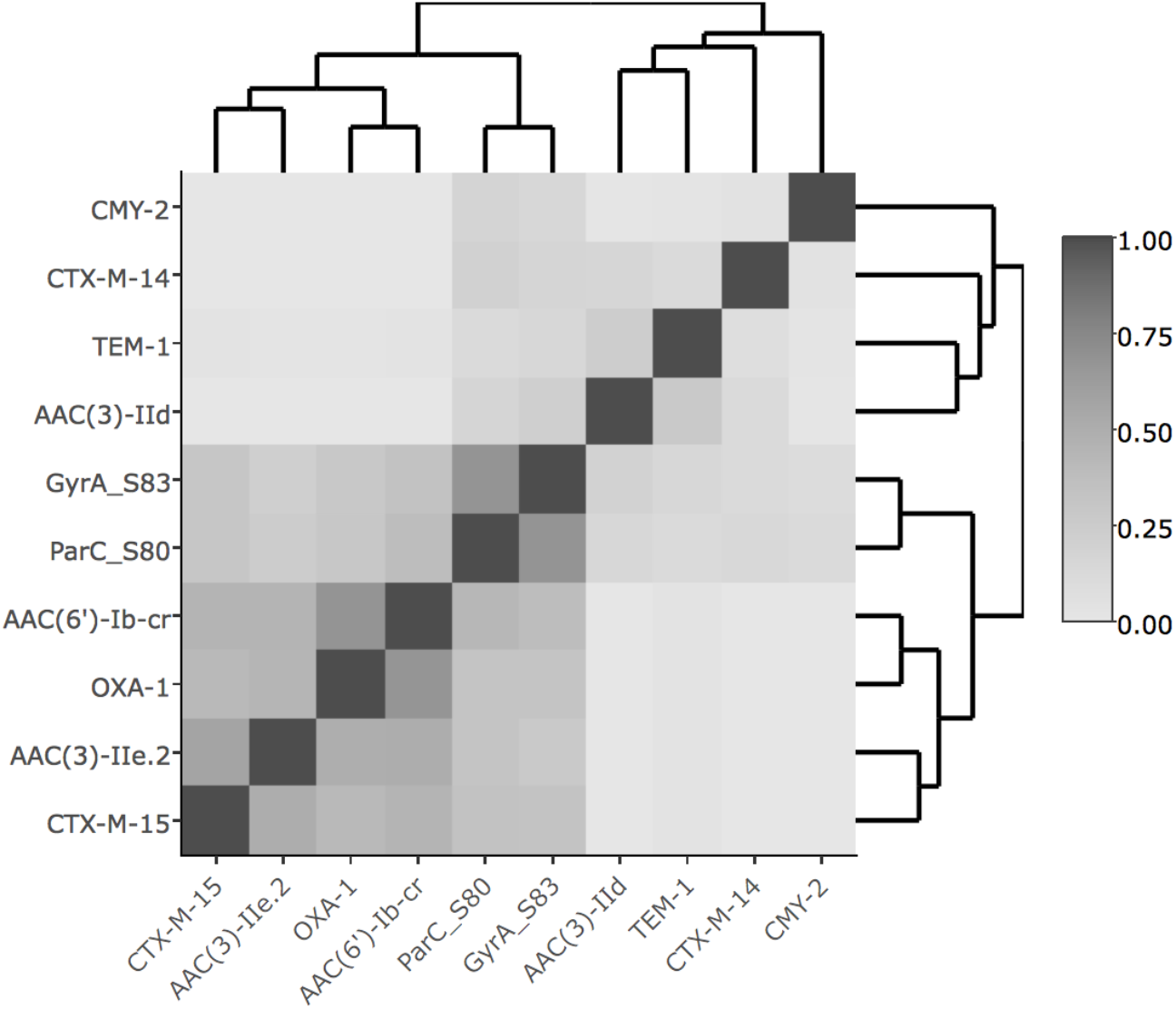
Pairwise associations between antibiotic resistance markers estimated based on their co-occurrence in genomes. The heatmap shows the strengths of associations calculated as log odds ratios normalized to between 0.00 (weak) and 1.00 (strong).

We also find that functionally redundant genes within these two groups rarely overlap. The two acetyltransferase genes showing strong correlations with gentamicin resistance (AAC(3)-IId and AAC(3)-IIe.2) do not co-occur in the 33 genomes containing these genes. The same was true when we compared the two acetyltransferase genes that correlated with resistance to tobramycin (AAC(3’)-IId and AAC(6’)-Ib-cr), which don’t co-occur in any of the 40 genomes where they were found. Similarly, we only found one case of co-occurrence between the two groups of ESBL *β*–lactamase genes seen in our samples: CTX-M-15 and CTX-M-14 family/CMY-2 in a total of 42 relevant genomes. In the case of OXA-1 and TEM-1 genes, the lack of overlap is not so sharp, but still substantial: out of 117 genomes containing TEM and 37 containing OXA, only 10 genomes have both.

In sum, we found that aminoglycoside and *β*–lactamase genes are genetically linked, although in some cases not very tightly, in two distinct clusters. We also see that when genes are functionally redundant, they rarely overlap. Therefore, the observed combinations of genes can be understood as two mutually exclusive complementarity groups: complementarity group 1 (CG1) and complementarity group 2 (CG2).

### Predictive value of individual genetic markers, alone and in combination

Next, we explored the predictive value of using antibiotic resistance genes, individually and in combination, as markers for epidemiological monitoring of resistance in the population and also for use in point-of-care diagnostics of antibiotic resistance. If two genes represent functionally redundant solutions for resistance to the same antibiotic, as our gene co-occurrence analysis suggests, each gene by itself should have low sensitivity, while including both genes should dramatically increase the sensitivity of our predictive models. Indeed, inclusion of both AAC(3)-IIe.2 and AAC(3)-IId for prediction of GEN resistance resulted in much greater sensitivity (82.5%) than when each was used as a marker individually (22.2% and 53.3%, respectively). Similarly, AAC(6’)-Ib-cr and AAC(3)-IId complement each other for prediction of TOB resistance, producing a sensitivity of 88.9%. For prediction of resistance against 3^rd^ and 4^th^ generation beta-lactams (CAZ, CRO, FEP), both CTX-M-15 and the CTX-M-14 family of beta-lactamases are needed to achieve high sensitivity (**Supplemental_Table_S2, Supplemental_Table_S3**).

### Mapping of CG1 and CG2 genes on our cladogram identifies multiple HGT events and allows inference of directionality

We next mapped our diagnostic CG1 genes encoding OXA-1, AAC(6’)-Ib-cr, CTX-M-15, and AAC(3)-IIe.2, and CG2 genes encoding TEM-1 and AAC(3)-IId on the phylogeny we generated based on SNP variation across the genome (**Fig. 1**). We hypothesized that when the host strains are ordered according to phylogenetic relatedness, the presence of these genes in non-contiguous host strains approximates independent occurrences. However, gene loss cannot be ruled out as an alternative explanation for the discontinuity of representation in the cladogram.

Our analysis identified 16 putative independent occurrences for CG1 (labeled 1 through 16 in **Table 1**) and 22 of them for CG2 (labeled A through V in **Table 2**). The OXA-1 and AAC(6’)-Ib-cr genes are found in all 16 putative occurrences, suggesting a strong genetic linkage between these two CG1 genes. Indeed, they are found adjacent to each other and flanked by insertion elements. CTX-M-15 is also almost always found as well. The exception is occurrence #7. Interestingly, in occurrence #2 CTX-M-15 is found in some samples but not all. Taken together, these observations suggest that CTX-M-15 is the third gene to be acquired in this group. AAC(3)-IIe.2 is found only in 9 of the 16 occurrences, and in occurrences with more than one sample (#3 and 9), it is not found in all samples. This suggests that AAC(3)-IIe.2 is the most recently acquired gene into this complementarity group.

**Table 1.**
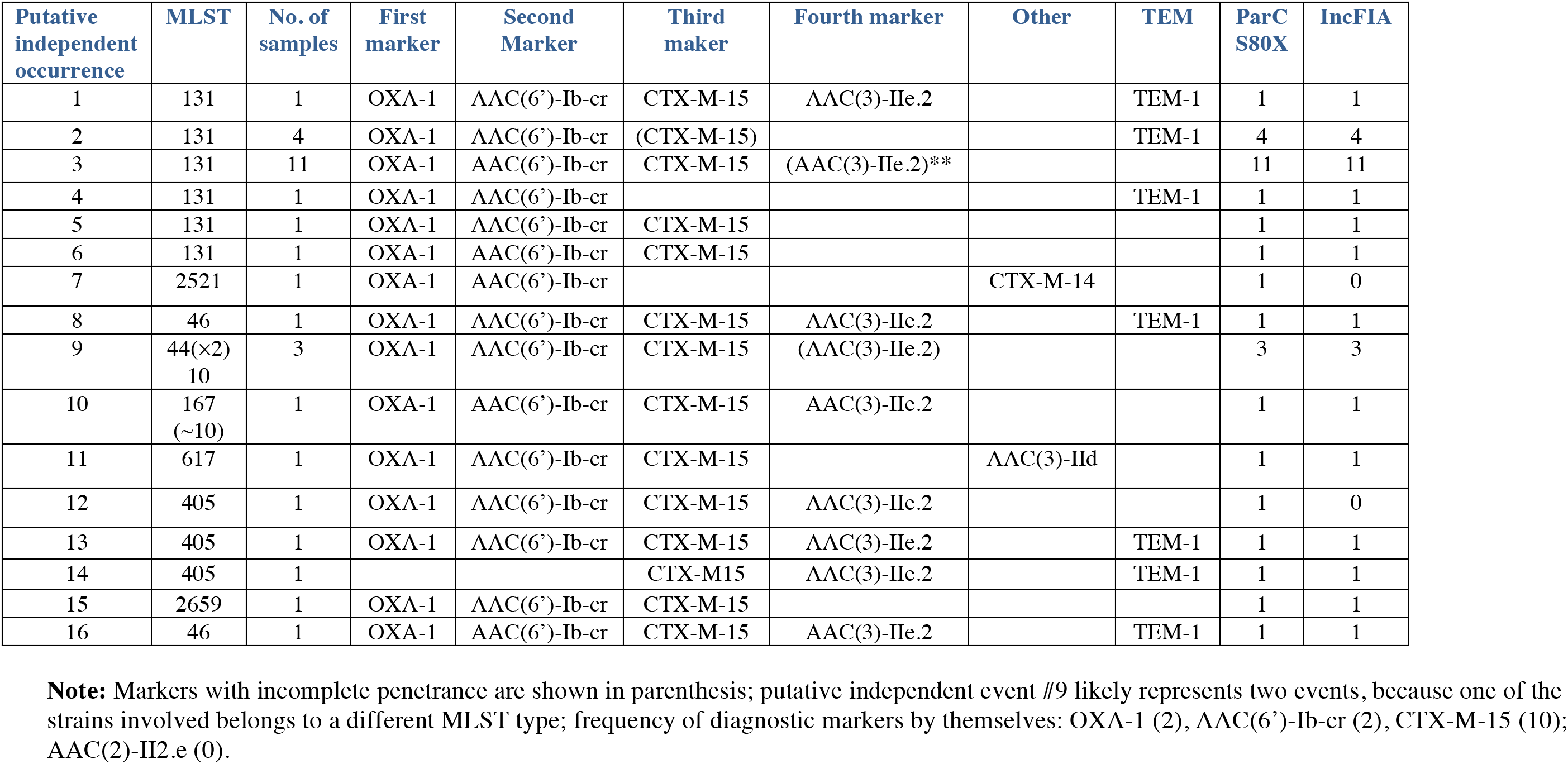
Complementarity group 1.

**Table 2.**
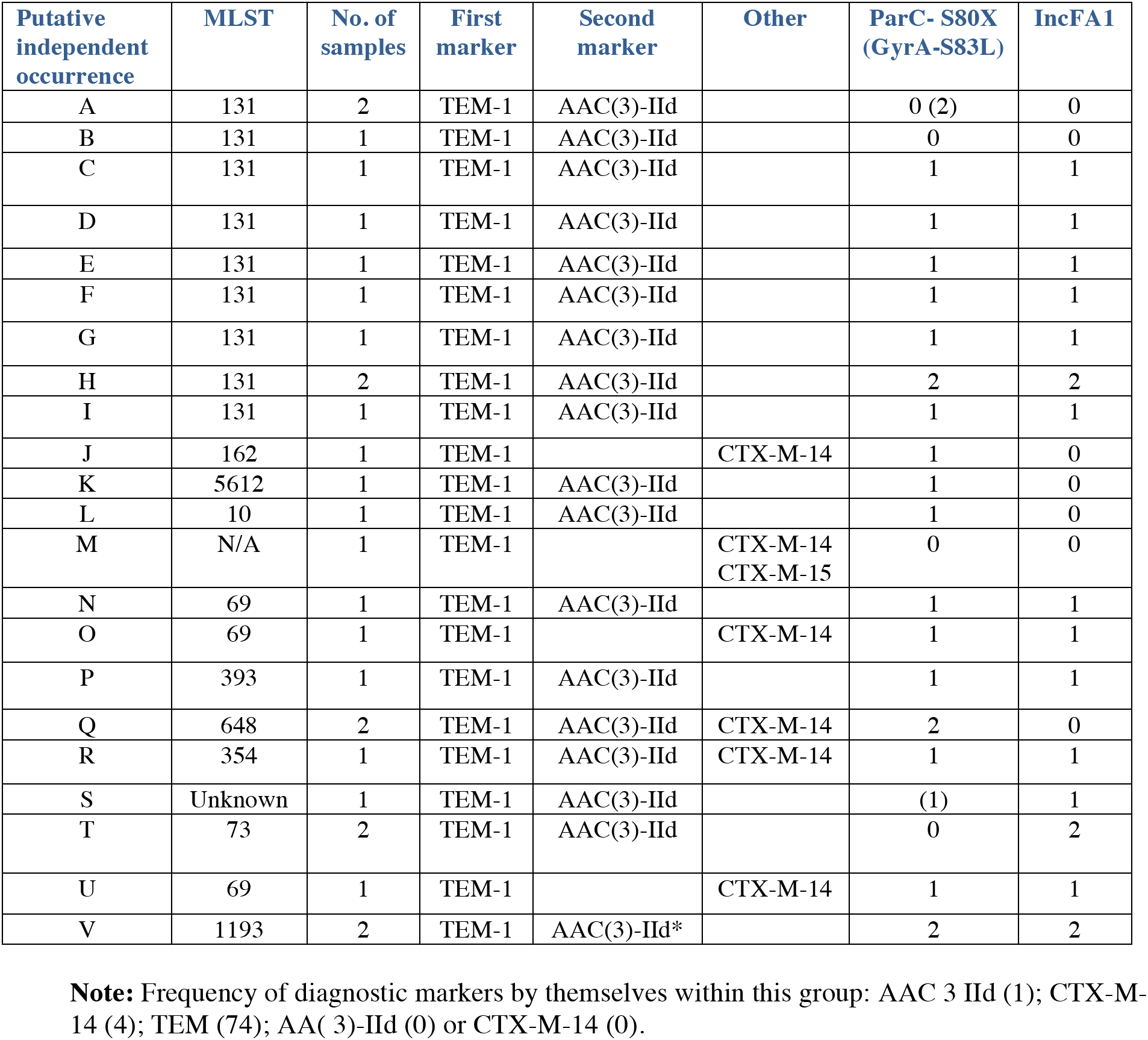
Complementarity group 2.

CG2 is characterized by the presence of AAC(3)-IId as opposed to other AAC enzymes, by a tight linkage to TEM-1 (100% of occurrences, compared to an average frequency of 44% in our population and in GC1 samples), and by a near complete absence of CTX-M-15.

### IncF plasmids are frequent and carry resistance to oxymino cephalosporins and aminoglycosides

IncF plasmids were very prevalent in our clinical population: 221 out of 276 strains carried an IncF plasmid, as determined by the presence of IncFIA, IncFIB or IncFII origins of replication.

CG1 and CG2 genes appear to be carried by IncF plasmids. In our data, aminoglycoside and ESBL resistance was found almost exclusively in the 221 genomes where IncF replicons were detected (with only 2 exceptions in the 55 remaining genomes). Further, IncF replicons were the only replicons consistently associated with AAC and plasmid-borne *β*–lactamase genes even though IncF-bearing genomes contained a variety of other replicons: IncI1, IncK, IncP, IncBO, IncAC, IncN, ColE1, IncR, IncY and IncLM (**Supplemental_Data_S1**).

The role of IncF plasmids in the acquisition of bla_TEM-1_ and bla_OXA-1_, in the subsequent capture of ESBL genes (Woodford et al. 2011; Olesen et al. 2013), and in the dissemination of CTX-M genes (Nicolas-Chanoine et al. 2008; Woodford et al. 2009; Hu et al. 2014) and of CMY (Matsumura et al. 2012; Naseer et al. 2012) by ST131 strains is well-established. Their role in spreading AAC(6’)-Ib-cr and CTX-M has been reported (Baudry et al. 2009; Li et al. 2015; Agyekum et al. 2016), and the AAC-(6’)-Ib-cr, OXA-1, and CTX-M15 combination has also been previously mapped to IncF plasmids (Li et al. 2015).

IncF plasmid origins of replication are modular (Hu et al. 2014). IncFII seems to be the essential element, present in nearly all plasmids. IncFIB is next in order of frequency (177), followed by IncFIA (106). **Fig. 3a** shows the average distribution of IncF origins of replication for our entire sample population. IncFIA is most frequently associated with both IncFII and IncFIB, whereas IncFIB is most frequently associated with IncFII alone, without IncFIA. This pattern of occurrence is consistent with a previous report (Hu et al. 2014) and suggests an asymmetrical dependence of IncFIA on IncFIB or a differential frequency of loss between the two subgroups.

**Figure 3.**
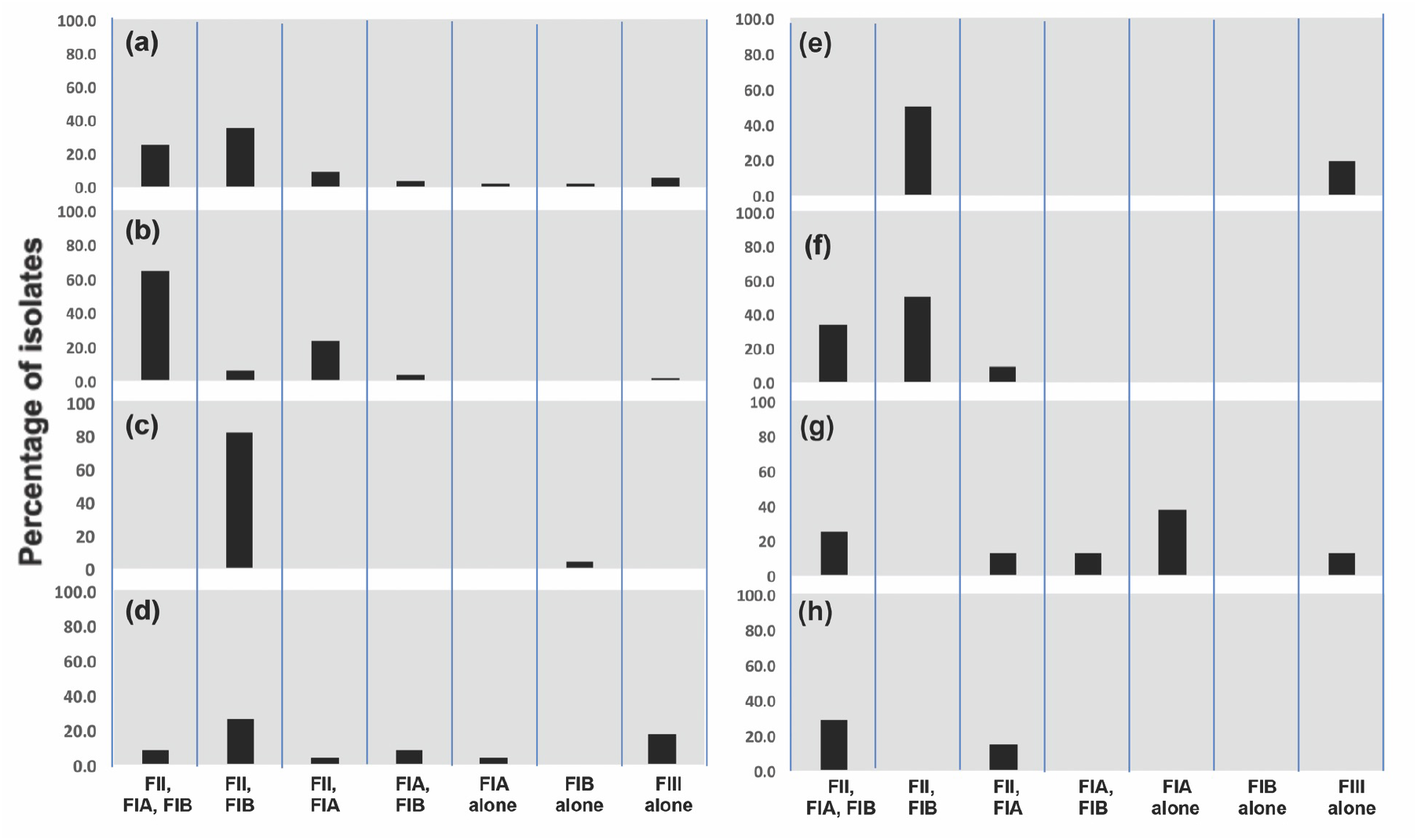
Distribution of IncF plasmid replicon type combinations across several prominent MLST groups. Proportion of isolates (y-axes) containing each combination of IncF replicon types is shown across the 276 ExPEC isolates, **(a)** as a reference, or in the samples belonging to the following MLST types: **(b)** ST131, **(c)** ST95, **(d)** ST173, **(e)** ST127, **(f)** ST69, **(g)** ST393 and **(h)** ST10.

**Fig. 3b-h** shows the average distribution of IncF origins of replication for the seven MLST types that are represented with at least 6 samples: ST131 (**panel b**), ST95 (**panel c**), ST173 (**panel d**), ST127 (**panel e**), ST69 (**panel f**), ST393 (**panel g**) and ST10 (**panel h**). The representation of IncFIA is strikingly variable depending on the MLST: from not represented at all (ST95, and ST127), to greatly overrepresented (ST131, ST10). IncFIA overrepresentation can be due to an increased representation of IncFIA alone (ST393) or of IncFIA in combination with IncFII and/or IncFIB (ST131, ST10).

### IncF plasmids form distinct clades

Next, we investigated the phylogenetic structure of the IncF plasmids present in our population. Given that plasmids are subject to frequent recombination and HGT, we focused our analysis on a single protein, the Rpt-A1 replicase, which is found in a region of the IncF plasmid that appears to be relatively stable (Villa et al. 2010; Shin et al. 2012). To further minimize noise from HGT and gene duplications, we limited our phylogeny to 204 samples carrying full-length (not truncated) Rpt-A1 sequences found as single copy genes (of 221 total samples carrying IncF plasmids). This analysis indicated the presence of seven distinct classes of IncF plasmids (**Fig. 4**). To reinforce our conclusions from our phylogenetic analysis of a single IncF plasmid protein (Rpt-A1), we performed Gaussian Mixture Modeling with Expectation Maximization to generate a second grouping of plasmid sequences based on the presence or absence of multiple IncF-specific sequence elements including various replication origins, replicase genes and relaxase genes. This analysis identified nine clusters (**Supplemental_Fig_S3**).

**Figure 4.**
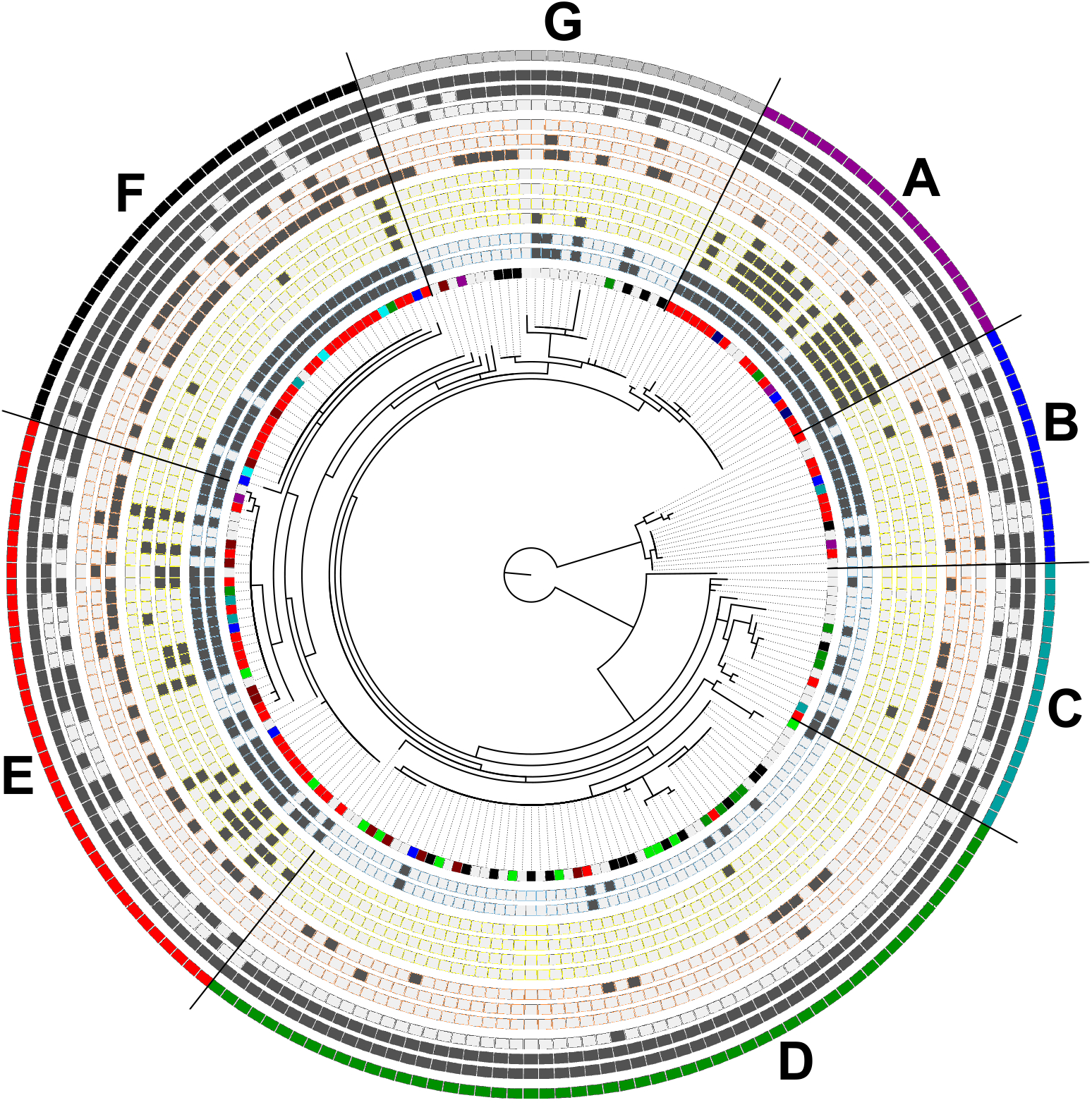
Maximum-likelihood phylogeny of 203 IncF-specific replicases, Rpt-A1. Genomic assemblies encoding multiple Rpt-Al of canonical length, but with diverged sequences, were not included in the phylogeny in order to minimize noise from HGT and gene duplications. The Circos diagram surrounding the phylogeny shows for each phylogenetic taxa represented by one unambiguous Rpt-Al sequence, its MLST classification, presence of various resistance markers and plasmid replicons. Circular panels represent the following, from the innermost to outermost position. **Panel 1**, MLST classification: ST131 (red), ST44, (navy), ST10 (purple), ST405 (blue), ST69 (green), ST393 (teal), ST648 (cyan), ST73 (maroon), ST95 (black), ST127 (lime). **Panels 2** and **3** (outlined in blue) fluoroquinolone resistance mutations: GyrA-S83X and ParC-S80I, respectively. **Panels 4-8** (outlined in yellow), complementarity group 1: genes for OXA-1, AAC(6’)-Ib-cr, CTX-M-15, and AAC(3)-II2.e, respectively. **Panels 9-11** (outlined in orange), complementarity group 2: genes for TEM-1, CTX-M-9-family and AAC(3)-IId, respectively.

**Panels 12-14** (outlined in grey), IncF replicons: IncFIA, IncFIB, IncFII, respectively. **Panel 15**, phylogenetic clades assigned manually upon careful inspection of Rpt-A1 phylogeny clade structure.

### Detection of IncF horizontal and vertical transfer

Given that we previously established that MLSTs closely approximate phylogenetic relatedness in our population (**Fig. 1**), we investigated whether the seven classes of plasmid defined above were restricted in their host strain distribution, focusing on MLSTs that are well represented in our population (with more than five samples). The number of different plasmid classes found in each of these MLSTs is shown in **Supplemental_Fig_S4a**. This number ranged from 2 to 6, which is very high considering that some of these MLSTs were represented by fewer than 10 genomes. The distribution of plasmid groupings obtained from our Gaussian Mixture Modeling analysis across MLSTs also suggested extensive conjugation among different plasmid groups (**Supplemental_Fig_S5**). The most parsimonious explanation for the presence of different classes of IncF plasmids in strains belonging to the same type is HGT by conjugation.

The presence of plasmids belonging to the same class in samples that are contiguous in the cladogram and that belong to the same MLST is suggestive of clonal expansion, because in this case vertical transmission of the plasmid from a common ancestor is the most parsimonious explanation. This analysis points again to the four frequent MLSTs identified earlier as likely representing recent clonal expansion, namely ST131, ST95, ST73, and ST127. In addition, this analysis found two additional, less frequent MLTSs (3 samples each) that we would not have otherwise been able to detect: ST648, and ST421.

### CG1 and CG2 are carried by distinct sets of IncF plasmids

We mapped the occurrences of the genes defined in CG1 and CG2 to our Rpt-A1-based phylogeny of IncF plasmids (**Fig. 4)** with the assumption that these genes are largely carried in IncF plasmids. The gene-to-isolate correspondence is provided in **Supplemental_Table_S4**. We found that each CG is carried by distinct sets of IncF plasmids.

CG1 is largely found in classes A and E. In class A, the linkage between genes encoding OXA-1, AAC(6’)-Ib-cr and CTX-M-15 is stronger than in class E (100% vs 50%). Class E also has a higher representation of CTX-M-14-like genes (4 samples, compared to 1 for Class A). Clade F has one instance of the complete set of CG1 genes including one encoding AAC(3)-Ile.2, suggesting that despite the inconsistent linkage between the genes that define CG1, they can be moved together.

For CG1, there are two additional associations: with fluoroquinolone resistance mutations, and with IncFIA. ParC-S80I/R is present in 79.6% of the samples, but in 100% of the samples carrying CG1 genes. Likewise, while IncFIA represents 72% of the total combining IncF plasmid classes A and E, it is present in 93% of the samples carrying CG1 genes. These two associations are specific for CG1: for CG2, ParC-S80I/R was found in 74% of the samples (77% expected combining classes B and F) and h IncF1A was found in 74% of the samples (67% expected).

CG2 corresponds to clades B and F in our plasmid phylogeny. Clade F has the same frequency of CTX-M-14 and of CTX-M-15, though.

Finally, we looked to see whether the different classes of IncF plasmids were differentially distributed in clonally expanding strains. **Supplemental_Fig_S5b** shows the breakdown of classes represented in our frequent MLSTs (black columns), compared to the distribution found in rare MLSTs (white columns). All seven classes have a good representation in both groups of strains, again consistent with ubiquitous HGT. We found that the classes associated with GC1 and CG2 drug resistance (A, B, E and F) are overrepresented in clonally expanding strains, whereas classes C and G (with a weaker association with drug resistance) are more frequent in rare strains. This seems to corroborate the idea that high risk clones help spread resistance.

## Discussion

Here we describe the presence of two sets of genes in clinical samples from two hospitals on the West Coast of the US: the first set (CG1) includes four genes encoding drug-inactivating enzymes: OXA-1, AAC(6’)-Ib-cr, CTX-M-15 and AAC(3)-IIe.2. The second set (CG2) includes genes for TEM-1, AAC(3)-IId and CTX-M-14-like enzymes.

Using linear regression analysis, we established strong correlations between the presence of these genes and resistance to some aminoglycosides and oxyimino-cephalosporins (**Supplemental_Fig_S2)**, and using these genes, constructed strong predictive models for these drugs (**Supplemental_Table_S2** and **Supplemental_Table_S3**). Tobramycin and ampicillin were exceptions, suggesting the presence of additional mechanisms of resistance to these drugs that we have not been able to identify.

The repeated identification of sequencing assemblies corresponding to a few genes, with conserved boundaries across different contexts (multiresistance regions or MRRs) has been described in the past (Partridge 2011). The genes belonging to our two CGs likely comprise MMRs, although we have not verified their location in the sequence relative to each other (with the exception of AAC(6’)-Ib-cr and OXA-1 which occur in close proximity, and bounded by conserved insertion elements). These CG genes do co-occur much more frequently than would be expected by chance (**Tables 1, 2** and **Fig. 1)** and parts of the two CGs that we describe here have already been reported as MRRs: the co-occurrence of AAC-6’-Ib-cr with CTX-M-15 (Gibreel et al. 2012; Banerjee et al. 2013), and the combination of AAC-6’-Ib-cr, OXA-1, and CTX-M-15 (Li et al. 2015).

Here we report three novel findings regarding the linkage between these resistance genes: (1) that CTX-M-14-like genes are found in a separate CG relative to CTX-M-15; (2) that a new AAC(3) aminoglycoside acetyltransferase closely related to AAC(3)-IIe (AAC(3)-IIe.2) is in the process of being acquired by CG1; and (3) that each CG is linked to a different *β*–lactamase gene: OXA-1 (CG1) and TEM-1 (CG2), although TEM-1 is ubiquitous.

We also show that these previously reported gene associations are very prevalent and explain the bulk of gentamicin and ESBL resistance cases in our two U.S. West Coast hospitals. Remarkably, this includes not only ST131 types but also many others (**Tables 1,2**).

ST131 is unique in that it is a carrier for genes belonging to both CGs with similar frequency: 10% of ST131 isolates carry CG1 genes and 14% carry CG2 genes. However, for the most part, CG1 and CG2 have different host types. Frequent CG1 carriers include ST44, ST46, and ST405; others include ST617, ST2521, and ST2659. By contrast, ST69 is the most frequent carrier for CG2 genes (25% of 12 isolates); other CG2 hosts include: ST73, ST354, ST393, ST1193 and ST5612. These findings extend previous reports indicating an association of specific MLSTs with CTX-M enzyme dissemination: ST131 and ST405 in the case of CTX-M-15 (Woodford et al. 2011; Canton et al. 2012; Shin et al. 2012), and ST393 in the case of CTX-M-14 (Canton et al. 2012). The multiplicity of MLSTs seen is consistent with the widespread mobilization of IncF plasmids across types shown in **Fig. 4**.

We find that when CG1 and CG2 genes are functionally redundant, they almost never overlap in the same isolate. This pattern suggests that the gene set fixed in a given population is largely the result of contingent evolution; once a strain acquires one MRR, an alternative MRR no longer provides a selective advantage. This is the reason why we propose calling these linked gene sets “complementarity groups”. The exceptions are OXA-1 and TEM-1, which in this case appear to follow a mutually-exclusive distribution pattern because of their linkage to AAC(6’)-Ib-cr (CG1) and AAC(3)-IId (CG2), respectively. Thus, in this case mutual exclusion is indirect, and therefore weaker.

However, the linkage between the genes in each complementarity group is dynamic, which allows us to infer the order in which the resistance markers are being acquired following the principle of genetic capitalism originally postulated by (Baquero 2004), **Fig. 1** and **Tables 1, 2**.

In the case of CG1, the initial mutations appear to be mutations in GyrA and ParC conferring fluoroquinolone resistance. This had been previously reported in the context of ST131 (Naseer and Sundsfjord 2011), but here we show that this phenomenon applies more broadly to strains harboring CG1 genes (**Table 1**, second column). The OXA-1 + AAC(6’)-Ib-cr combination is the next one to appear; the two genes are adjacent and flanked by insertion sequences, which explains their tight linkage. The next gene to be captured is CTX-M-15, almost certainly as a result of treatment with oxymino cephalosporins or with aztreonam. Finally, the last gene, which is very much in the process of capture into CG1 is AAC(3)-IIe.2, likely as a result of gentamicin selection.

CG2 appears to have arisen by the capture of AAC(3)-IId, linked to TEM-1. Subclass II AAC(3) acetyltransferases are common and confer resistance to both tobramycin and gentamicin (Shaw et al. 1993), consistent with our linear regression analysis, which suggests that this gene provides resistance to both tobramycin and gentamicin (**Supplemental_Fig_S2**). The next gene, which is in the process of capture is CTX-M-14 in the broad sense (also including other enzymes of the CTX-M-9 cluster).

The apparent connection between fluoroquinolone resistance (which is chromosomal) and the IncF-borne CG1 aminoglycoside and ESBL resistance genes is not obvious and intriguing. It could simply be the result of the timing of selection, with fluoroquinolones being used as first line of defense and ESBLs as an alternative when fluoroquinolones fail, but this does not explain why this pattern is not seen for the CG2 group. Alternatively, AAC(6’)-Ib-cr, which confers fluoroquinolone resistance, may be selected under fluoroquinolone selective pressure as an enhancer of the resistance provided by chromosomal mutations in GyrA and ParC. This would explain, why a separate AAC (AAC(3)-IIe.2) gene is selected later on to provide gentamicin resistance. This would also explain why in strains harboring CG2 genes, where no specific linkage to fluoroquinolone resistance can be detected above a high background of resistance, a single gene that provides resistance to both gentamycin and tobramycin was captured.

We identified a few sequence types that are undergoing clonal expansion and that therefore constitute potential high-risk clones (Woodford et al. 2011): ST131, ST95, ST73, ST127, and ST69. This conclusion is based on representation (all have at least 12 samples), on a short phylogenetic distance between samples belonging to the same type, and on evidence of vertical transmission of CG genes (since this only happens following acquisition of this putative MRR and is therefore a relatively recent evolutionary event). Further, we observed that ST131 strains were much more abundant in the DHMMC samples (which were obtained more recently), representing 67.4% of the total, compared to 16.5% in the UW study (which was 2–3 years older), consistent with the idea that ST131 strains continue to expand in the clinical population. We confirmed that this apparent overrepresentation of ST131 strains in the DHMMC set cannot be attributed to its different antibiotic resistance profile, by performing Monte Carlo sampling distributions matching the proportions of ceftazidime-, ciprofloxacin- or gentamycin-resistant samples among the 43 DHMMC assemblies (see *Methods, Supplemental Materials* and **Supplemental_Fig_S6**).

The ST131 type is known to have undergone a dramatic expansion since they were first reported in all three continents in 2008 (Simner et al. 2011; Colpan et al. 2013). It is interesting to note that the three high-risk clones with the highest representation in our study (ST131, ST73 and ST95) all belong to the phylogroup B2 (Le Gall et al. 2007), suggesting that this expansion is not simply driven by selection for drug resistance (many other STs also have these genes) but by some ecological/virulence advantage. There is strong evidence that this is the case for ST131 strains, whose expansion is at least partially driven by the acquisition of FimH, a mannose-binding type 1 fimbrial adhesin, which appears to facilitate uroepithelial colonization by *E. coli* (Johnson et al. 2013). On the other hand, we did find IncF plasmid classes associated with CG1 and CG2 drug resistance (A, B, E and F) are overrepresented in clonally-expanding strains, supporting a role for these strains in the spread of antibiotic resistance. The fact that only a subset of plasmids belonging to these classes carry CG1 or CG2 genes suggests that they are facilitating the spread of resistance but not necessarily driving it.

IncF plasmid replicons are modular and complex, frequently including multiple typing subgroups (Hu et al. 2014). Similar to other studies (Villa et al. 2010), we find that IncFII is the essential one, frequently associated with IncFIB, which in turn can be associated with IncFIA(**Fig. 1, 4**). Here we show that IncF replicon modules can be highly MSLT-specific (**Fig. 3**).

We also show an association of IncFIA with the plasmid classes bearing CG1 and CG2 genes (frequency of IncFIA representation range of 60–95%, compared to 2–35% for the other plasmids); in classes A and E, there was an additional association between CG1 resistance genes and IncFIA: 93% of strains carrying CG1 genes also carried IncFIA origins of replication, compared to 72% of stains carrying class A and E plasmids. These observations suggest a preferential role of IncFIA origins of replication in the mobilization of antibiotic resistance.

Our Rpt-A1 phylogeny also found 13 instances of likely co-occurrence of IncF plasmids, based on the identification of two Rpt-A1 copies in the same sample (**Supplemental_Table_S5**). In all but one case, the two copies belonged to a different plasmid class, indicating that these do not represent recent duplication events. Interestingly, in all but two cases, one of the plasmids belonged to class B, the one class that is clearly more phylogenetically different from the other six (**Fig. 4**). This observation suggests that class B plasmids may be frequently compatible with other IncF classes. The observation that IncF plasmid co-occurrence appears to depend on the phylogenetic distance of Rpt-A1 suggests that compatibility is not a process primarily driven by the IncF subgroup complement. Instead, the modularity of IncF plasmid replicons may represent an adaptation to the host rather. This would be consistent with the observation of IncF signatures for individual MLST types reported here.

## Concluding remarks

Here we describe a binary gene distribution of *β*–lactamase and aminoglycoside resistance genes in the clinical population of two West-Coast hospitals. This observation suggests an evolutionary convergence between different evolutionary solutions, producing ubiquitous functional redundancy. The discovery of specific complementarity groups and the functional redundancy they represent has significant implications for monitoring and controlling the spread of drug resistance. For example, functional redundancy greatly complicates the rational design of antibiotic therapeutic regimens aimed at maintaining the population sensitive (Mira et al. 2015).

We were able to build models with high predictive accuracies (90–95%) for most drugs using combinations of only 2–3 genes. Sequence-based predictions offer three critical advantages over clinical microbiology assays (1) reproducibility, *i.e*. by using a standardized analytic pipeline, results can be reproduced which is important for clinical-related usage, (2) traceable historical data, which helps monitor the spread of resistance, and (3) contextual information, which allows estimation on the likelihood of resistance development to additional antibiotics [reviewed in (Didelot et al. 2012; Burnham et al. 2017)]. However, the extent to which the markers that we have identified can be generalized to other clinical populations is unclear.

Accurate prediction of resistance phenotype at point of care would help delay the spread of resistance, minimizing misuse of antibiotics, which currently increases both the costs of treating infection and accelerates the spread of resistance. On the other hand, evidence of pervasive HGT and recombination indicates a great potential for the acquisition and spread of new resistance genes that would need to be incorporated to our predictive models if these are to remain accurate.

## Acknowledgements

We would like to acknowledge Dr. Steve Salipante’s help processing and interpreting the UW sequences, Dr. Gerardo Cortés-Cortés for useful comments on the manuscript. This work was funded by CITRIS Seed Funding proposal 2015–324 to TL, MC and MB, and by NIH/NIAID award 1R41AI122740–01A1 to PC, MC and MB.

## Methods

### Sample and data collection

Clinical samples were collected from patients with respiratory, blood (wound), or urinary tract infections at Dignity Health Mercy Medical Center (DHMMC) in Merced, California, between June 2013 and August 2015. The isolates were identified as ESBL-positive using an automated rapid detection system for pathogen identification and antibiotic sensitivity, Vitek 2 Version 06.01. Following identification, the samples were also tested for susceptibility against 16 antibiotics using broth micro-dilution minimum inhibitory concentration (MIC) testing. The isolates were categorized according to their susceptibility: Resistant (R), Intermediate (I), or Susceptible (S), based on the MIC Interpretation Guideline – CLSI M100-S26 (2015). The 16 antibiotics included 1 penicillin: Ampicillin, 2 penicillin and inhibitor combinations: Ampicillin/Sulbactam, Piperacillin/Tazobactam, 4 cephalosporins: Cefazolin, Ceftazidime, Ceftriaxone, Cefepime, 2 carbapenems: Ertapenem, Imipenem, 3 aminoglycosides: Amikacin, Gentamycin, Tobramycin, 2 fluoroquinolones: Ciprofloxacin, Levofloxacin, and Nitrofurantoin and Trimetroprim/Sulfamethoxazole.

Additionally, we obtained 342 ExPEC genome assemblies from a previous study conducted at the University of Washington (UW), downloaded from Genbank (Benson et al. 2016) with accessions in the range JSFQ00000000–JSST00000000.

### Sequencing, quality control and assembly of ExPEC genomes

Genomic DNA was extracted from each sample using the ZR-96 Quick-gDNA Kit from Zymo Research. Whole genome sequencing including TruSeq DNA library preparation was conducted at the University of California, Davis Genome Center using Illumina’s MiSeq and HiSeq technologies. We obtained 24 MiSeq (2×250 bp) and 24 HiSeq (2×250 bp) paired-end sequencing libraries corresponding to our selected samples. Prior to assembly, Illumina sequencing adapters and low quality bases were trimmed from the sequencing reads using Trimmomatic v0.36 (Bolger et al. 2014); trimmed reads shorter than 36 bp were discarded. Library quality was also verified using FastQC v0.11.5. *De novo* paired-end assembly was conducted for each MiSeq and HiSeq library using SPAdes v3.5.0 (Bankevich et al. 2012) with read error correction by BWA-spades. The spades.py wrapper script was used to select an appropriate *k-* mer size for optimized assembly of each genome.

The resulting assemblies were scanned for contamination and misassembly; contigs shorter than 300 bp or with low coverage were discarded. Estimated genome sizes (median: 5.3 Mb) based on the remaining contigs were generally within the range of known ExPEC genomes. The assemblies had 29x and 327x median coverage for MiSeq and HiSeq libraries, respectively. The median N50 across 22 MiSeq and 23 HiSeq library assemblies was 198 kb (median L50: 9) and 201 kb (median L50: 8.5). Two HiSeq libraries appeared to contain significant contamination or proportions of misassembled contigs based on estimated genome sizes (8.2 Mb and 9.0 Mb for samples 328 and 357, respectively); these two assemblies were removed from downstream analyses. We further assessed each DHMMC and UW assemblies for completeness based on the presence of 143 protein-coding genes considered to be essential for normal growth of *E. coli* MG1655 [Set A in Supplementary Table 2 from (Gerdes et al. 2003)]. To achieve consistent assembly qualities across the DHMMC and UW datasets, we removed assemblies containing fewer than 126 full-length essential genes.

### Phylogeny and typing of strains

We conducted multi-locus sequence typing (MLST) of the 276 genome assemblies based on 15 housekeeping loci required for categorizing *E. coli* strains according to the Achtman (7 loci) and Pasteur (8 loci) MLST schemes.

To estimate phylogenetic relationships among the 276 assemblies, we constructed a neighbor-joining tree based on all possible pair-wise comparisons. For each pairing, an evolutionary distance based on single nucleotide polymorphisms (SNPs) was estimated using the nesoni package [ref]. SNPs for individual assemblies were detected by aligning all sequencing reads comprising an assembly against an *E. coli* reference genome, EC958, which belongs to the ST131 (Forde et al. 2014). Note that a phylogeny generated in this manner may be subject to noise arising from SNPs found among horizontally transferred elements (e.g., genes in plasmids and genomic islands). However, we believe such noise should have little influence on the shape of the phylogeny since the number of SNPs found in vertically transmitted regions should greatly outnumber the SNPs found in horizontally transmitted regions. We confirmed the general accuracy of our SNP-based phylogeny by comparison against a maximum-likelihood phylogeny generated using the concatenated protein sequences of 126 essential *E. coli* genes.

A second maximum-likelihood phylogeny based on Rpt-A1 protein sequence homology was generated to approximate the evolutionary relatedness of IncF plasmids. To minimize noise from HGT and gene duplications, we first removed assemblies putatively encoding multiple, different Rpt-A1 proteins of canonical length (285 amino acids) *i.e*. assemblies with multiple Rpt-A1 were included only if there was 100% sequence agreement. Maximum-likelihood phylogenies were generated using RAxML v8.2.10 (Stamatakis 2014); bootstrapping was performed on the CIPRES compute cluster (Miller et al. 2015). Multiple alignments for these trees were generated using MAFFT v7.294b (Katoh and Standley 2013); gaps were removed using TrimAl (Capella-Gutierrez et al. 2009).

### Gene model prediction and annotation

Gene models were predicted using Prodigal v2.6.2 (Hyatt et al. 2010) in metagenomic mode. Known antibiotic resistance-conferring genes and “mutations” were annotated using NCBI BLASTx and BLASTp (Altschul et al. 1990) based on amino acid residue-level differences between selected “wild-type” genes and full-length hits (greater than or equal to 99% query coverage and greater than 90 percent identity). Selected “wild-type” genes included known resistance genes encoding β-lactamases (TEM-1, SHV-1, OXA-1, CTX-M-14, CTX-M-15), aminoglycoside-modifying enzymes (acetyltransferases [AAC], adenyltransferases [ANT], phosphoryltransferases [APH]), 30S ribosomal protein subunit S12, 16S rRNA methyltransferases (ArmA, RmtA, RmtB, RmtC, RmtD, RmtE), DNA gyrase subunits (GyrA and GyrB), DNA topoisomerase IV subunits (ParC and ParE), and plasmid-mediated quinolone resistance proteins (QnrA, QnrB, QnrB2, QnrC, QnrD, QnrS). Amino acid sequences and Uniprot (UniProt 2015) accessions for wild-type genes are provided in **Supplemental_Data_S2**. Plasmid replication origins and associated incompatibility groups were annotated based on BLASTn hits (e-value < 10^-50^) against INC-DB (Lanza et al. 2014), a plasmid origin nucleotide sequence database.

### Predictive values and co-occurrence of known genes

The predictive values of individual genes and combinations of genes were assessed based on Pearson correlation and several other measures including sensitivity, specificity, positive predictive value (PPV) and negative predictive value (NPV). The relative importance of each marker for resistance prediction was estimated as regression coefficients obtained from a generalized logistic regression prediction model, fitted using the glm package in R. Reported coefficients were the average result of 1000 bootstrap iterations. Marker co-occurrence associations were assessed using adjusted p-values and log odds ratios calculated using Fisher’s exact tests.

## Abbreviations

ESBL: Extended Spectrum Beta-Lactamase
CG: complementarity group
HGT: horizontal gene transfer
Inc: plasmid incompatibility
AAC: aminoglycoside acetyl transferase

